# Mechanistic Insight from Full Quantum Mechanical Modeling: Laccase as a Detoxifier of Aflatoxins

**DOI:** 10.1101/2020.12.31.424992

**Authors:** Marco Zaccaria, William Dawson, Darius Russell Kish, Massimo Reverberi, Maria Carmela Bonaccorsi di Patti, Marek Domin, Viviana Cristiglio, Luca Dellafiora, Frank Gabel, Takahito Nakajima, Luigi Genovese, Babak Momeni

## Abstract

We demonstrate a path towards full Quantum Mechanics (QM) characterization of enzymatic activity. As a case-study, we investigate the detoxification of aflatoxin, a carcinogenic food contaminant, by laccase, a versatile oxidase capable of—but not efficient for—degrading aflatoxin. We use a combination of quantitative experimentation and QM modeling to show that low enzymatic steric affinity for aflatoxin is the main bottleneck, rather that the oxidative activity of laccase. To identify the structural elements responsible for low reaction rates, we perform a density functional theory (DFT) based modeling of both the substrate and the enzyme in a full QM simulation of more than 7,000 atoms. Thanks to our approach we point to amino acid residues that determine the affinity of laccase for aflatoxin. We show that these residues are substrate-dependent, making a full QM approach necessary for enzyme optimization. Altogether, we establish a roadmap for rational enzyme engineering applicable beyond our case study.

## Introduction

State-of-the-art in-silico characterization of enzyme-substrate systems relies on hybrid quantum mechanical/molecular mechanical (QM/MM) simulation methods. QM/MM requires preparatory set-up to identify an apt region for the QM simulation, while the rest of the enzymatic structure is modeled with a more computationally manageable MM description. Determining the appropriate QM region is a crucial and delicate step in QM/MM models [1]. QM regions including ≥6 amino acid residues (about 100 atoms) are conventionally regarded as large. In this work, we implement an approach to include a complete protein-substrate system in a full QM simulation. Our model, encompassing all laccase’s amino acid residues plus the aflatoxin molecule, quantum mechanically simulates approximately 7600 atoms. This methodological development is implemented using the BigDFT code [2].

Achieving full QM characterization of enzyme-substrate interactions would allow an agnostic approach to enzymatic characterization, eliminating the need for prior knowledge about relevant QM regions in the QM/MM model [3], [4]. It would also allow validation of previous QM/MM models, enable detailed characterizations of less studied/tractable enzyme-substrate systems, and reveal suitable simplifying assumptions. The required CPU demands for such a computational paradigm have, until recently, made it unapproachable at the relevant dimensional scale. For a first attempt at a full QM model, an experimentally tractable enzyme-substrate system which enabled reliable model verification would be a promising candidate. We introduce the laccase-aflatoxin interaction as such a system. The motivation behind this is multi-faceted: (i) laccase is an enzyme of general interest in biotechnology for which mechanistic enzyme characterization and optimization would be beneficial [5], [6]; (ii) in the context of bioremediation, the current state-of-the-art is inadequate for targeting aflatoxins—among the most dangerous food pollutants [6]; and (iii), current QM/MM modelling appears to be incapable of explaining the laccase-aflatoxin experimental results (Supplementary Information).

Laccase is a monomeric multicopper oxidase: it catalyzes one-electron oxidation of its substrate coupled with full reduction of molecular oxygen to water. The active site is a functional unit formed by three types of copper binding sites with different spectroscopic and functional properties. Type 1 blue copper is the primary electron acceptor from the substrate, while a trinuclear cluster formed by type 2 copper and binuclear type 3 copper constitutes the oxygen binding and reduction site [7]. Laccase is taxonomically ubiquitous [8] and functionally versatile: its broad substrate tolerance enables it to catalyze a range of oxidative reactions relevant to several industrial applications [5]. Across natural variants, fungal laccases have the highest redox potential (E°)—up to 800 mV, at the type 1 copper [8]. However, even the strongest natural isoforms cannot offer a time/cost efficient aflatoxin bioremediation process. Characterizing the mechanisms behind detoxification could realistically enable functional optimization, but metallo-proteins such as laccase are particularly challenging for QM/MM formalizations.

Mycotoxins are dangerous fungal secondary metabolites that regularly contaminate staple crops such as maize, small cereals, rice, and peanuts [9]. The most dangerous mycotoxins are the aflatoxins, produced by *Aspergillus* species, also arguably the most carcinogenic natural occurring pollutants [10]. Aflatoxin contamination is a major food safety concern. Physical and chemical aflatoxin decontamination strategies exist, but they are costly, unsafe, or with undesired side effects [11], [12]. As a promising alternative, food recovery through environmentally safe enzymes has been proposed [13]. To this end, laccase has been identified as a good candidate [14]–[16].

Our goal is to identify the intrinsic limitations of laccase as an aflatoxin detoxifier to devise strategies to overcome them. In this research, particular attention is given to the detoxification of aflatoxin B_1_ (AFB_1_), the most carcinogenic of the aflatoxins. For detoxification experiments, we employed the laccase from the basidiomycete *Trametes versicolor* (TV), a fungal species whose ecological niche is tailored around laccase-mediated lignin degradation [17]. We perform an in-depth analysis of the detoxification of the main target molecule, AFB_1_, by TV laccase to identify the mechanisms behind reaction bottlenecks. Our data also include experiments on an AFB_1_ congener, aflatoxin G_2_ (AFG_2_), to compare and contrast our findings.

We first constructed a preliminary phenomenological model based on laccase activity on aflatoxins *in vitro*. The model highlighted two salient points: (i) laccase’s efficacy against aflatoxin is not limited by the redox potential of its active site, rather by poor affinity for aflatoxin as a substrate; (ii) AFB_1_, unlike AFG_2_, deviates from the established Michaelis-Menten kinetics characteristic of laccase activity [18]. Thus, improving the affinity appears to be the best option to achieve successful large-scale application. At the same time, remarkable differences in the detoxification of AFB_1_ and AFG_2_, despite their structural similarity, imply that affinity improvement would not be achieved only by specializing the enzyme towards a general category of compounds (e.g. hydrocarbons, aromatic nonphenolic structures, or even aflatoxins as a category). Instead, it would require a highly detailed approach that is capable of distinguishing between different congeners; to this end, we proceeded to perform quantum mechanical (QM) characterization of select variables of the laccase-aflatoxin system.

We simulated the effect of a laccase-like single-electron oxidation of aflatoxins using QM density functional theory (DFT) calculations of the neutral and oxidized molecules. In particular, we examined the Fukui functions—describing changes in the electron density in response to electron loss—to understand what parts of toxin molecules might get involved during toxin oxidation. Our results suggest the following scenario: (i) laccase does not achieve aflatoxin detoxification through a one-electron oxidation alone, and an ulterior environmental stimulation is required for detoxification (i.e. lactone ring opening); and (ii) the observed discrepancies in AFB_1_ versus AFG_2_ detoxification can be partially attributed to the intrinsic properties of the two toxin variants.

Moving forward, we turned to a full *ab initio,* DFT-based QM model of enzyme-substrate interaction. By including laccase in the QM model, we showed that we are able to (i) identify the amino acid residues that are pivotal to detoxification for the two tested aflatoxin variants; (ii) categorize the identified residues based on their degree of enzyme-substrate influence; and (iii) evaluate quantitatively how interactions between laccase and the two aflatoxin variants are variant-specific in terms of their strength and localization. We conclude that rational engineering of a laccase-based aflatoxin bioremediator mandates a detailed mechanistic approach. We present a systematic way to inform rational enzyme specialization to target a specific substrate. This approach is general and can be expanded to other enzyme-substrate pairs of interest.

## Results

### Laccase is a more effective detoxifier of AFG_2_ than it is of AFB_1_

In the chemical structure of aflatoxin, the lactone ring is responsible both for toxicity [19] and the natural fluorescence of the molecule. As a result, aflatoxin concentration and toxicity can be fluorimetrically assayed. In this work, we will define detoxification as a reaction that breaks the lactone ring in the aromatic structure of aflatoxin, leading to loss of natural fluorescence and toxicity [19]. We can thus correlate successful aflatoxin detoxification to loss of native fluorescence. This assay can be used for both AFB_1_ and AFG_2_ (see Methods).

To assess detoxification, we tested 50 U/mL of *T. versicolor* laccase at different initial AFB_1_ and AFG_2_ concentrations. The fluorimetric assay highlights two distinct detoxification trends for AFB_1_ and AFG_2_ by laccase. AFB_1_ fluorescence readout follows a decreasing trend that, after about 10 hours, changes into a slower trend. Overall detoxification over 96 hours is about 12% of the original quantity of the toxin (Fig. 1A). AFG_2_ detoxification, in contrast, displays a consistent trend, leading to complete detoxification of AFG_2_ within 96 hours (Fig. 1B).

**Figure 1.**
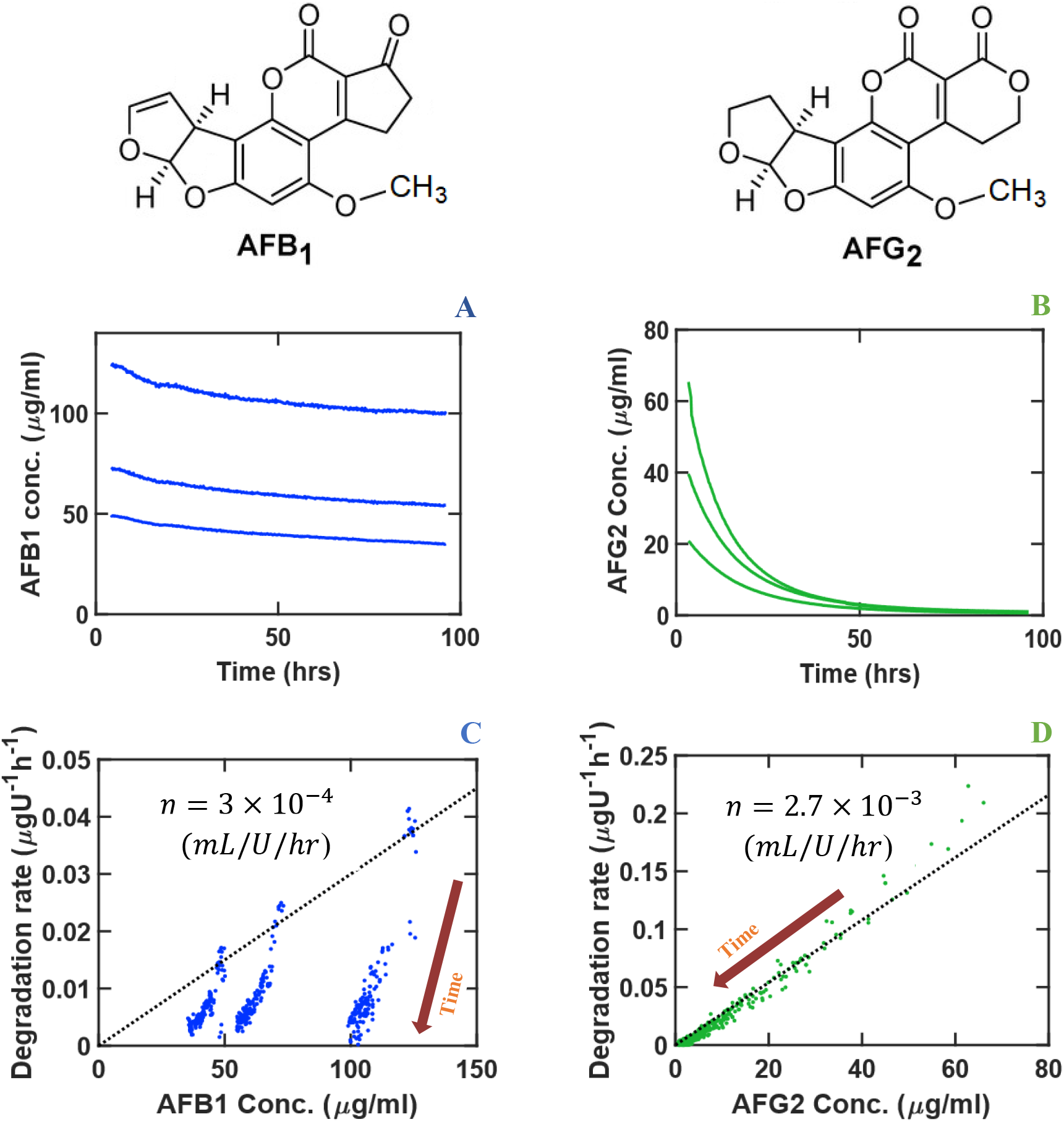
The detoxification of AFB_1_ and AFG_2_ by laccase highlights the difference in detoxification efficiencies even between aflatoxins with similar structure. Different initial aflatoxin concentrations were employed and are represented for AFB_1_ (A) and AFG_2_ (B). Each curve is the average of 3 replicates. A subset of points from (A) and (B) is randomly selected and represented in (C) and (D) to calculate the local normalized detoxification rates 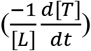. Detoxification efficiency 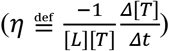 of AFB_1_ (C) is almost an order of magnitude lower than that of and AFG_2_ (D) at comparable concentrations. Dotted lines in (C) and (D) illustrate the prediction of the model, assuming *K_m_* ≫ [*T*] (see Equations 1 and 2). Direction of time is represented in (C) and (D) to highlight the decrease in toxin concentration as a result of detoxification. Laccase concentration: 50 U/mL.

### Laccase has higher affinity and a higher detoxification rate for AFG_2_ over AFB_1_

To infer the enzymatic activity of laccase against aflatoxins, we fitted a phenomenological model into our experimental data. In our model, we assumed that laccase detoxifies aflatoxins following the Michaelis-Menten equation:

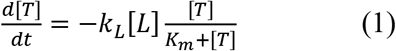

in which [*T*] is the toxin concentration (in 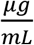), [*L*] is the laccase concentration (in 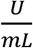), *K_m_* is the Michaelis-Menten constant (in 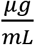), and *k_L_* is the degradation rate by laccase from the enzyme-toxin associated state (in 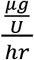).

In the limit that the toxin concentration is much lower than the Michaelis-Menten constant (*K_m_* ≫ [*T*]), the equation will be simplified to

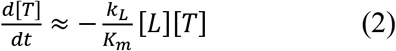

To test if this assumption is valid, from the experimental data we define detoxification efficiency 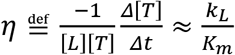 (in *mL*/*U*/*hr*), where *Δ*[*T*] is the change in the toxin concentration in a small time-step *Δt*. Since we can measure [*T*] experimentally over time, we can calculate *η* as well as the local normalized detoxification rate 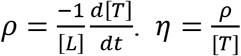 appears to be constant in early degradation (i.e. a linear trend in Figs. 1C-D), suggesting that *K_m_* ≫ [*T*] is a valid approximation. Calculating the detoxification kinetics from Eq (2), and using the value of *η* estimated from experimental data, the model accurately approximates the measured kinetics in the case of AFG_2_ throughout the experimental time (Fig. 1D), further confirming that this model is suitable for representing aflatoxin detoxification by laccase. However, AFB_1_ adheres to the Michaelis-Menten kinetics only for a short time before entering a slower, non-Michaelis-Menten-like detoxification dynamic (Fig. 1C). Thus, compared to AFG_2_, and other known substrates of laccase [18], AFB_1_ reacts uncharacteristically.

The finding that, at relevant concentrations of the toxin, we get *K_m_* » [*T*] can be interpreted as relatively poor activity by laccase for degrading the toxin. If we consider the association and enzymatic activity in the standard view [20]:

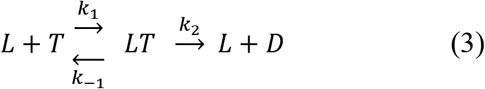

where *D* is the detoxified toxin, and 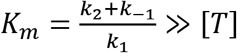 means *k*_2_ + *k*_–1_ ≫ *k*_1_[*T*]. This can be interpreted as low affinity of the enzyme for the aflatoxin, AFB_1_ and AFG_2_ alike, as the rate of association is much smaller than the rates of degradation/dissociation. This low affinity suggests that laccase is naturally not well-adapted to detoxify aflatoxin. As a matter of fact, it has been reported that under optimized conditions (0.1 M citrate buffer pH 4.5, 20% DMSO 35°C, TV laccase 30 U/mL) *K_m_* for AFB_1_ was 0.28 mM and degradation rate with 80 μg/mL AFB_1_ was *k_L_* = 0.89 μg/U/day [21]. This corresponds to a detoxification efficiency of *η* = 4 × 10^-4^ *mL/U/hr*, which is comparable to our results reported in Fig. 1C.

We make two main observations:

1. <u>Differences in enzyme-substrate affinity between AFB_1_ and AFG_2_</u>. *η* represents the detoxification efficiency and describes how the substrate is affected by the active site and consequential lactone ring opening. For AFB_1_, *η* = 3×10^-4^ *mL/U/hr;* in the case of AFG_2_, *η* = 2.7×10^-3^ *mL/U/hr*. Thus, laccase is an order of magnitude less efficient at detoxifying AFB_1_ compared to AFG_2_.
2. <u>Differences in detoxification kinetics.</u> AFG_2_ detoxification rate proceeds uniformly over time, whereas AFB_1_ detoxification becomes considerably less efficient after about 10 hours. This suggests a different mode of interaction between the enzyme and AFB_1_ versus AFG_2_.

### The multiplicity of oxidation sites may explain some of the differences between AFB_1_ and AFG_2_ detoxification

To investigate the differences observed between AFB_1_ and AFG_2_, we investigated the properties of these molecules in the gas phase using a QM model (see Methods for details). The isosurfaces of the Fukui functions highlighted the sites involved in a hypothetical one-electron oxidation such as the one performed by laccase (Fig. 2). For both molecules, the model revealed that the oxidation sites are neither on the lactone ring nor on its immediate proximities. Thus, oxidation occurs away from the lactone ring itself. Nonetheless, the lactone ring opens up after AFB_1_ and AFG_2_ are exposed to laccase, as can be inferred by the loss of toxin fluorescence. We also experimentally confirmed this by LC-MS analysis of the degradation products (Fig 2-FS1 and Fig 2-FS2). Other oxidation products include the well-known epoxy- and dihydroxylated forms on the terminal furan ring (Fig 2-FS1 and Fig 2-FS2). However, the cationic molecules still display an intact ring and remain structurally stable in the QM model. Oxidation and detoxification therefore do not coincide, although they are interdependent events. The model also revealed that the AFG_2_ molecule possesses a more delocalized Fukui function than AFB_1_ and is therefore susceptible to oxidation of a larger number of sites than AFB_1_. This aspect could partially explain the differences in the *η* values between AFG_2_ and AFB_1_ in our earlier results.

**Figure 2.**
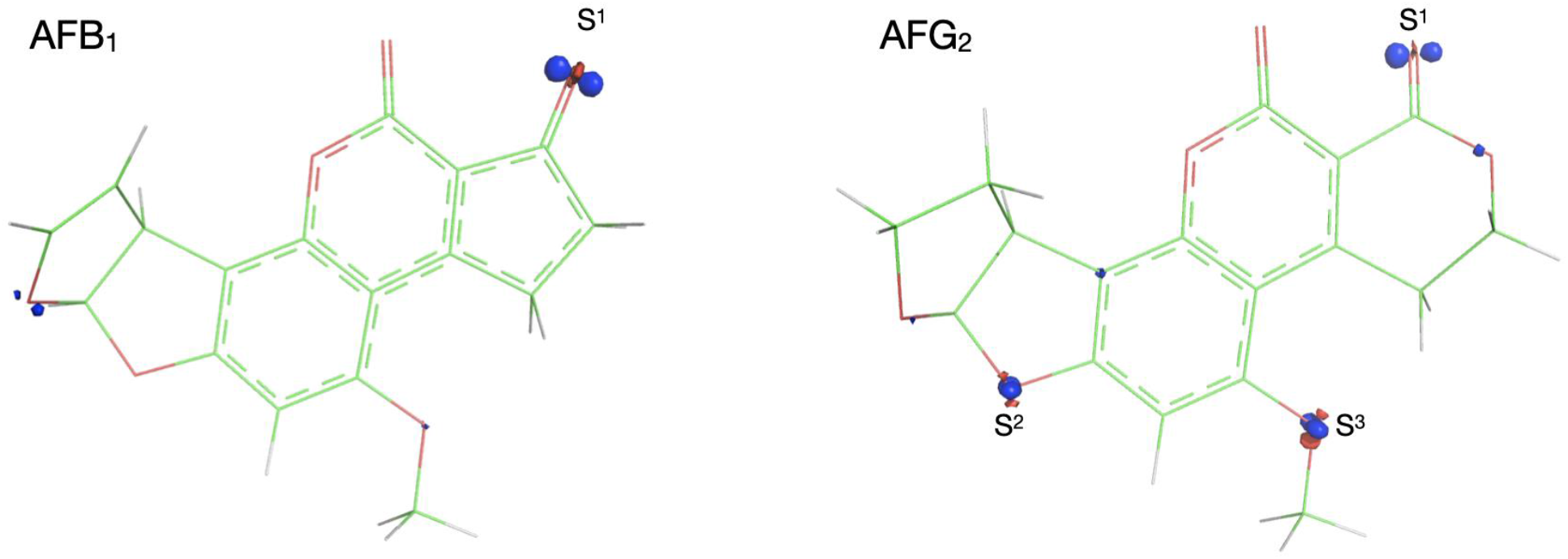
Isosurfaces of the Fukui functions of AFB_1_ and AFG_2_ in the gas phase indicate the sites prone to oxidation. Fukui isosurfaces: RED (-) and BLUE (+). In both molecules, oxidation occurs away from the lactone ring. AFG_2_ has a more delocalized Fukui function and is spatially more prone to oxidation.

In the QM model, we saw that the lactone ring did not spontaneously open after the toxin was oxidized in the absence of an environmental stimulation. Once we added an environmental stimulation in the simplest form of a hydrogen radical, the toxin spontaneously underwent a structural rearrangement that led to the formation of an epoxide in the terminal ring. We observed that when such environmental stimulation was localized in the immediate proximity of the lactone ring, the structural rearrangement would cause ring-opening (Fig. 3) without the need to overcome a barrier. AFG_2_ exhibited a lower free-energy conformation post-ring breakdown (−1.71 eV compared to oxidized state) compared to AFB_1_ (−1.34 eV compared to the oxidized state), which suggests a slightly higher tendency towards this transition.

**Figure 3.**
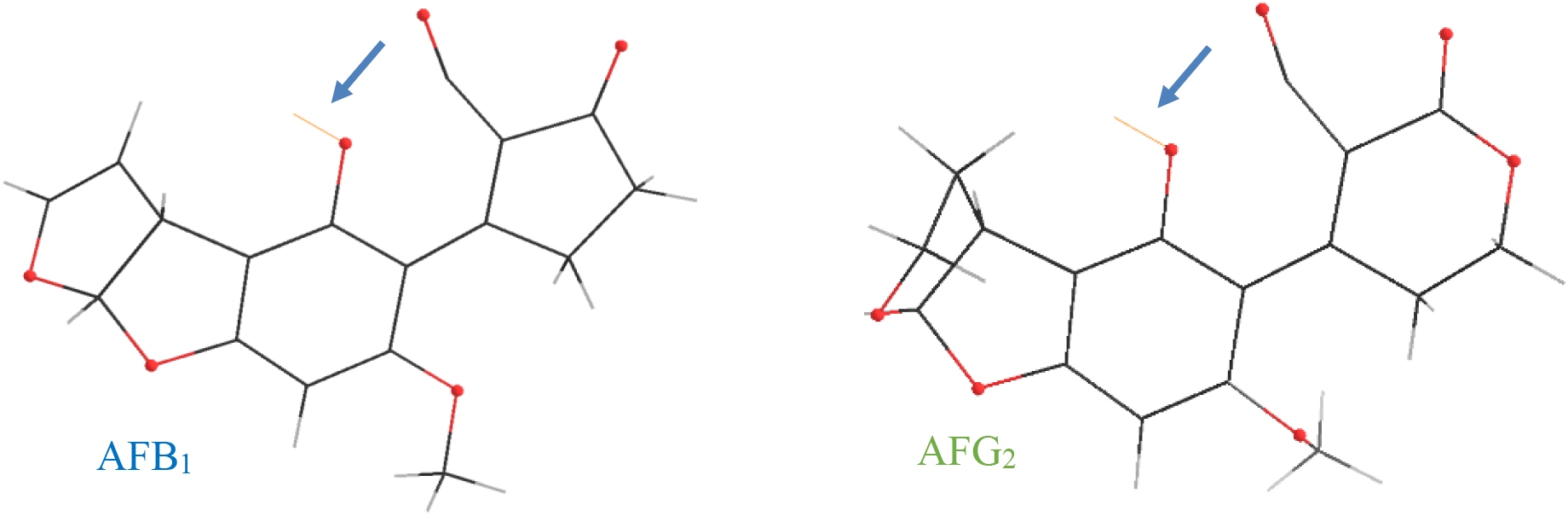
Conformations of the AFB_1_ and AFG_2_ during the simulated lactone-ring opening process indicate the position of environmental stimulation. We present a snapshot of the geometry optimization procedure of the oxidized aflatoxin + H system, showing how this would lead to a ring opening. The attacking hydrogen is shown by the yellow bond (marked by the arrow).

### Analysis of the binding site shows substrate-specific interaction of aflatoxins with laccase

To further study the difference in reaction dynamics between the two toxins, we used QM modeling of the docked toxin-enzyme system. We generated a set of docking poses for AFB_1_ and AFG_2_ in complex with the Beta isoform of laccase, as well as ABTS—a compound routinely used to measure laccase activity [22]—as a reference. From the set of computed docking poses, we selected five different geometries for each system since docking scores cannot reliably predict binding positions. We then performed geometry optimization on each system using a force-field model followed by QM calculations of the full system (Methods).

In our earlier publications [2], [23], we described a *Complexity Reduction* framework which uses the electronic density computed by QM calculations to reproduce the properties of a full system from calculations on only a subset of the system. The key ingredient of this analysis is the *Fragment Bond Order,* which is a generalization of atomic bond order to interactions between two arbitrary sets of atoms. The *embedding environment* for a given fragment contains all its major inter-fragment interactions and is defined as the minimal set of fragments such that the sum of the bond orders of all excluded fragments is below some threshold. Here, we propose to use this framework to generate interaction maps between the bound toxin and the different amino acids of laccase. This approach enables us to define a substrate-specific active site of laccase.

In Fig. 4, we show a graph view of these interactions for each of the different geometries. To generate this view, we computed the union of the embedding environments for a given substrate and the lone copper atom using a cutoff of 0.001. In our previous publication [23], we showed that properties converge with a cutoff between 0.01 and 0.001. Amino acids are colored by the cutoff required to add them to the embedding environment. Images of each of the substrate-interactors using the same color scheme is included in the Supplementary Information.

**Figure 4.**
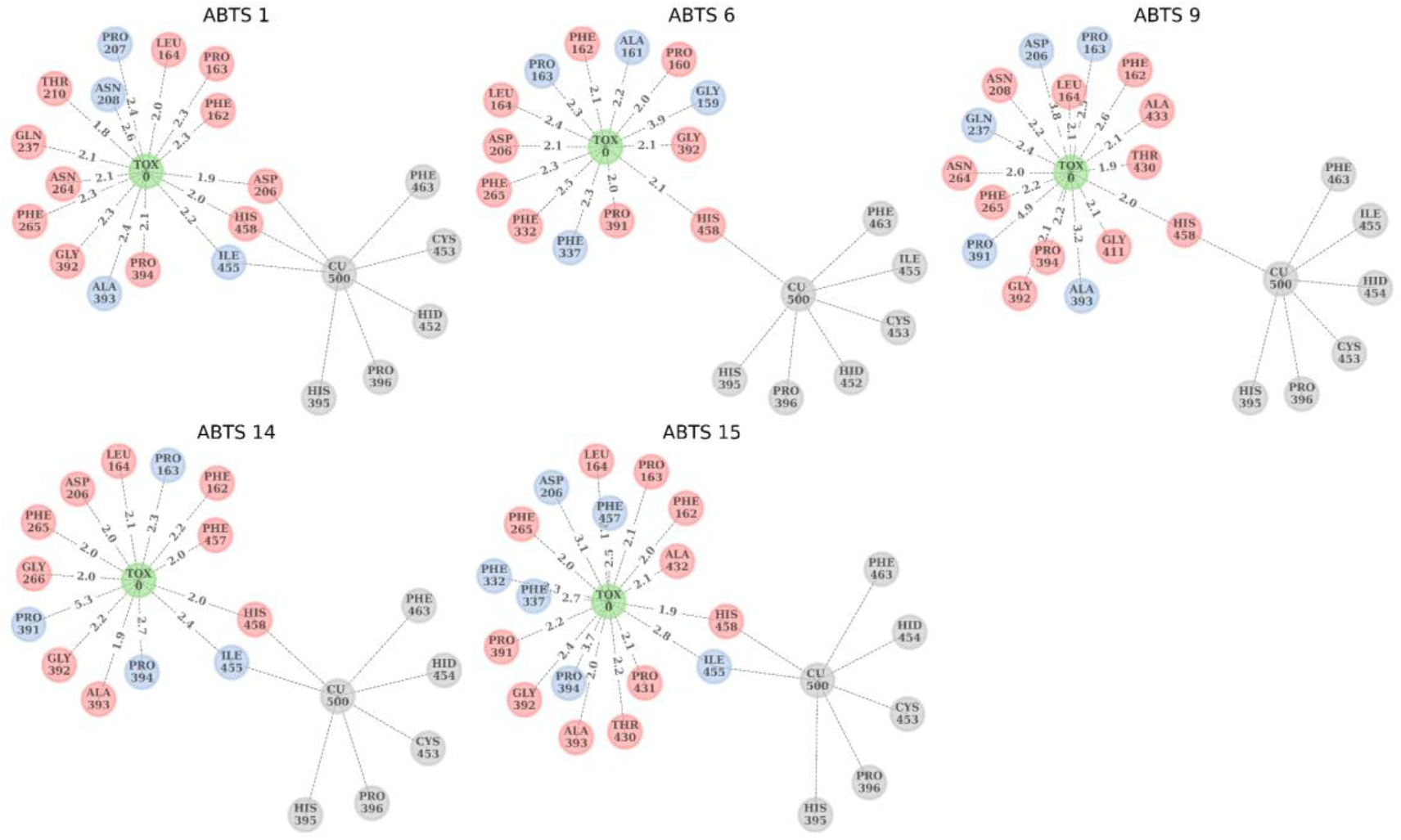
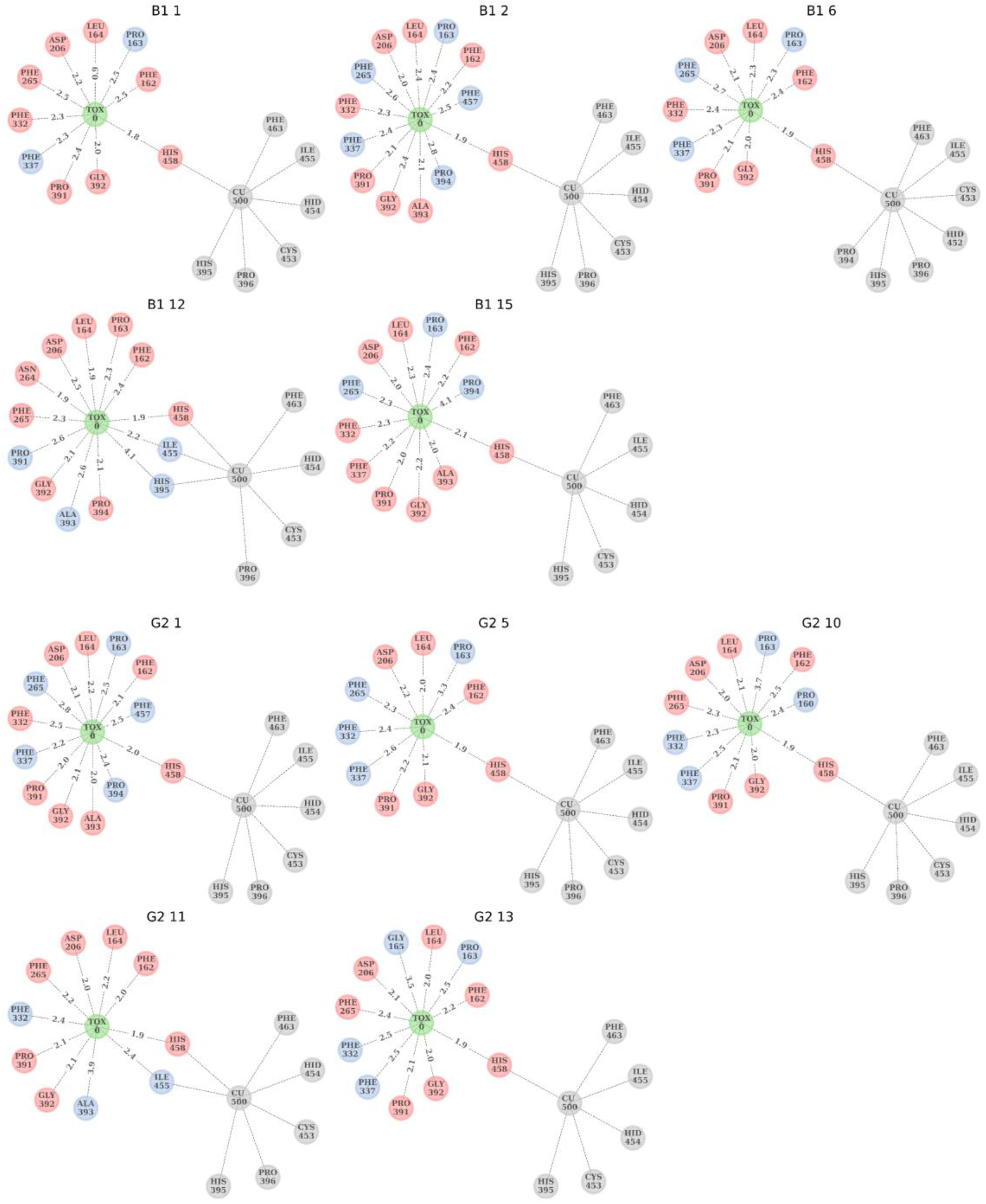
Complexity Reduction framework reveals that interactors are substrate-specific. The substrate is colored in green and labeled as TOX0. Amino acids that would be included with a cutoff 0.01 are colored in red and 0.001 in blue. Fragments that interact with copper, but not with the substrate are in gray. The copper is labeled as CU500. The edges of the graph label the nearest neighbor distance between each amino acid and the substrate (Å). Representative geometries of substrate-laccase structure are shown in Fig 4-FS1.

One trend we observe is the strong interaction of both the substrates and the copper atom with amino acid His458 of the catalytic site. This suggests that the distance between a substrate and this amino acid controls the activity of laccase. This finding echoes earlier studies [24], [25] which have suggested that substrates should dock within 5 Å of that residue. We also see interactions between Ile455 and His395 from the evolutionary preserved binding pocket, but this interaction is not consistent across poses. In Table 1, we report the distance of each system and their nearest Fukui function from His458. For ABTS, we also found that most poses lead to a Fukui function close to His458. However, we found that the Fukui function site shared by AFB_1_ and AFG_2_ is consistently too far away for efficient degradation. AFG_2_ has a unique Fukui site that is a similar distance to His458 as ABTS, a fact that may account for the differences in degradation rate between AFB_1_ and AFG_2_ despite their structural similarity. The analysis performed here suggests that a significant redesign of this laccase’s pocket that causes the shared Fukui function to be oriented inwards will be required to improve degradation efficiency.

**Table 1.** Calculated GOLDScores and Fukui distances are listed for different poses of ABTS, AFB_1_ and AFG_2_. GOLDScores represent the quality of docking. Fukui distance shows how far the potential oxidation site in the Fukui function is from His458 in laccase’s enzymatic pocket.

| System | GOLDScore | Fukui Distance (Å) |
| --- | --- | --- |
| ABTS 1 | 58.34 | 1.98 |
| ABTS 6 | 54.81 | 8.03 |
| ABTS 9 | 50.02 | 1.99 |
| ABTS 14 | 43.85 | 2.03 |
| ABTS 15 | 41.73 | 1.93 |
| B1 1 | 50.34 | 10.09 |
| B1 2 | 49.31 | 6.31 |
| B1 6 | 48.51 | 9.96 |
| B1 12 | 46.22 | 10.45 |
| B1 15 | 42.86 | 7.39 |
| G2 1 | 50.57 | 5.43 |
| G2 5 | 49.54 | 1.87 |
| G2 10 | 46.86 | 1.92 |
| G2 11 | 45.94 | 1.88 |
| G2 13 | 45.19 | 1.89 |

We caution that a more realistic set of atomic positions than those presented here would be required to fully understand the effects presented here. Using Mulliken population analysis, it is possible to compute the degree of oxidation of the substrate in each pose. We find that none of the substrates have been oxidized, which makes it clear that each substrate must first enter into an oxidized transition state before degradation occurs. Furthermore, the positions here are only based on relaxed crystal structure geometries and do not account for the full dynamics of the system in solution (see Discussion).

## Discussion

Until recently, technical limitations have prevented full QM *in silico* characterization of biological systems with hundreds to thousands of atoms, including enzyme-substrate complexes [26]. A formalized mechanistic description of such complexes would greatly benefit directed enzyme evolution and molecular engineering efforts for diverse applications (e.g. vaccines, aptamers, antibiotics, and bioremediators). We offer a basis to this approach, focusing on laccase-aflatoxin a tractable enzyme-substrate system of health relevance.

We have chosen the laccase-aflatoxin system as a case-study, because of laccase’s importance in biotechnology and aflatoxin’s health-relevance. Aflatoxin’s potential threat is highlighted in the recent alert issued by the U.S. Foods and Drugs Administration on contaminated pet food [27]. Aflatoxin contamination is indeed a major concern among food-safety issues, and laccase-mediated detoxification can be viewed as a promising “green” bioremediation approach [8]. The evolution of this lignin-degrading enzyme has led to an active site with the highest redox potential among multi-copper oxidases [5]. Such uncommon oxidative potential is a necessary asset for breaking down the aromatic moieties of aflatoxins. However, as our data highlight, laccase is not optimized to carry out this reaction and, not surprisingly, lacks high affinity towards AFB_1_. Therefore, the biocatalyst needs to be improved before realistic application can be realized. To this end, the QM description of the enzyme-substrate complex can identify critical molecular determinants of catalytic activity, guiding the protein engineering efforts.

Our work addresses the aflatoxin contamination issue from a novel angle: a bottom-up understanding of the laccase-mediated detoxification process. We started by identifying the relevant variables in the dynamics of the laccase-aflatoxin interaction. In our experiments, a simple model of enzymatic detoxification highlighted how enzyme abundance, laccase oxidative power, and substrate affinity were sufficient to describe the dynamics of detoxification. Moreover, the model suggested that affinity and detoxification rate bear the same weight in the overall detoxification function: i.e., when contributing to the overall detoxification, a fold increase in affinity is equivalent to the same fold-increase in rate (Equation 2). Importantly, the laccase detoxification of AFG_2_ was much faster than AFB_1_ (Fig 1), although still far from the efficacy necessary to warrant realistic applications, with low affinity remaining the primary target for improvement. For bioremediation to have a realistic chance, it has to be implemented during the current food production process without disrupting it. With this in mind, the best context for laccase-mediated aflatoxin bioremediation would be during the water washing step in the production process of food commodities. For that, AFB_1_ detoxification would need to be achieved in no longer than about 3 hours, in a slightly acidic (pH of 6.5), aerobic, liquid environment at room temperature. Our data indicate that, at pH 6.5, even the detoxification of AFG_2_ by TV laccase takes >48 hours, far from what would be practically required.

The difference between detoxification of AFB_1_ and AFG_2_ is substantial and surprising (Fig 1), if we consider how structurally similar the two substrates are. These observations suggest that, to optimize laccase detoxification of AFB_1_, a more detailed study would not be an affectation, but a necessity. This necessity can be best addressed through a mechanistic, fully quantum mechanics-based approach. We find that higher level methods used previously, namely docking studies [28]–[30], were unable to elucidate the difference in binding observed. They are, however, still a necessary step when bound crystal structures are unavailable in order to generate a finite number of reasonable binding poses to be used in full QM studies. Our docking results, both GOLDScores and R-NiB rescoring [31] (see Supplementary Materials) showed little correlation with Fukui distance, and ranking between scoring methods were inconsistent with observed experimental ranking of performance. Although in our case docking alone was unable to elucidate differences in binding that explain differences in enzymatic degradation rate, it was necessary to generate poses for the QM studies that were consistent with these differences.

Our QM description provides three conclusions: (i) the lactone ring cannot open unless oxidation happens first; (ii) additional environmental stimulation at the lactone ring of the oxidized toxin is required to cause ring opening, i.e., laccase does not directly interact with the lactone ring to achieve detoxification and does not need to, because oxidation is achieved elsewhere; (iii) one possible explanation for the laccase’s higher affinity for AFG_2_ over AFB_1_ can be offered by the more delocalized Fukui function of AFG_2_, which makes AFG_2_ prone to oxidation from more than one site. This last point suggests that aflatoxins potential for detoxification by natural laccase may be limited, because it depends on the toxin’s intrinsic traits. Nonetheless, before ruling out the application of laccase for aflatoxin detoxification, the mechanistic details of the enzyme-substrate complex dynamics need to be formalized.

QM modeling is by definition a mechanistic approach and can be used to validate any alternative empirical modeling approach. Reservations about its employment are only limited to its feasibility: present-day computational power cannot reliably model QM-described atomic interactions of more than a few hundred atoms (when the dynamics of the atoms is considered). Simplifying the system to gas-phase QM calculations of the neutral and oxidized toxin may only provide speculative results of the actual processes eventually leading to detoxification. Recently, it has been shown that with a sufficient set of descriptors from docking and using quantum mechanical modeling of the gas phase substrate, it is possible to predict laccase affinity on a wide class of systems [25]. The specific case shown here, however, demonstrates a need for the combined modeling of the enzyme and substrate in order to properly predict detoxification. In this view, computational protocols like the QM/MM techniques (as employed for laccase in [32], [33]) may be employed. However, such techniques are based on chemical intuition for preliminary identification of the active site region which may be altered and modified by the actual conformation of the enzyme-substrate complex, as we have demonstrated in this paper for the particular case of AFB_1_ and AFG_2_.

The striking difference between the detoxification rates of AFB_1_ and AFG_2_ suggests that selecting for general categories of substrates would likely not be enough. Additionally, our data indicates that simpler models, such as molecular docking, are inadequate for providing the level of detail required for assessing detoxification efficiencies (e.g. discriminating AFB_1_ versus AFG_2_ responses). Our analysis has provided a map of the residues that play a direct role in the affinity of the toxin molecule for the enzymatic pocket, and also allowed an estimate of their intensity of interaction. In doing so, the QM model of the laccase-aflatoxin complex can inform efficient directed enzyme evolution.

A full QM approach could shed light on the following relevant questions: (i) Is the oxidation rate of the toxin dependent on how the latter approaches the active site? (ii) Is the enzyme active site altered by the toxin presence? (iii) How accurate are the QM/MM approximations with respect to an unbiased, full QM calculation? To answer such questions, we need a QM approach that is capable of handling systems with hundreds to thousands of atoms. The BigDFT code, employed in this paper for QM modeling, has been proven to be able to tackle Kohn-Sham Density Functional Theory (KS-DFT) calculations of systems up to a few tens of thousands of atoms [34], [35]. The resulting simulations can provide reliable information on the identification of the systems’ fragments and associated physical observables [23], [36].

The connection between the experimentally measured enzymatic efficiency and the mechanistic modeling depends on the reliability of the atomic coordinates employed for the *ab initio* modeling. As stated above, the mechanistic model we have considered for the docking is based on the laccase crystal structure, as detailed in a previous study [15]. However, it is important to assess the actual conformation that the enzyme may have in solution. To address this point, ongoing studies employ methods such as Small Angle X-ray Scattering (SAXS) and Small Angle Neutron Scattering (SANS) [45,46] to obtain low-resolution structural information of biological samples in solution.

To summarize, we provide a methodology for a full QM approach that can reveal the mechanisms of ligand-receptor interaction. Our observations can guide rational engineering efforts by providing insights about the ideal enzyme structure. For detoxifying aflatoxins in particular, we envision that such mechanistic modeling paves the way for *in silico* generation of theoretical laccase structures capable of efficiently detoxifying aflatoxin and also highlight laccase’s intrinsic limitations. The theoretical cues could then be evaluated experimentally by generating the respective protein mutants using site-directed mutagenesis.

Full QM characterization of other prototypical enzyme-substrate complex conformations using the methodology presented in this paper is ongoing. We envision that future work in this direction will drive rational engineering and directed evolution efforts for the development of efficient vaccines, aptamers, antibiotics, and bioremediators.

## Acknowledgements

We thank Laura Ratcliff, Claudia Mondelli, and Nicolas Coquelle for useful discussions. BM and MZ were supported by a start-up fund and an Ignite internal grant from Boston College and by an Award for Biomedical Excellence from the Smith Family Foundation. This work was supported by the Next-Generation Supercomputer project (the K computer) and the FLAGSHIP2020 project (Supercomputer Fugaku) within the priority study 5 (Development of new fundamental technologies for high-efficiency energy creation, conversion/storage and use) from the Ministry of Education, Culture, Sports, Science and Technology (MEXT) of Japan. Experiments presented in this paper were carried out using the Grid’5000 testbed, supported by a scientific interest group hosted by Inria and including CNRS, RENATER and several Universities as well as other organizations (see https://www.grid5000.fr). Additional calculations were performed using the Hokusai supercomputer system at RIKEN. This work was partly supported by Cabinet Office, Government of Japan, Cross-ministerial Strategic Innovation Promotion Program (SIP), “Technologies for Smart Bio-industry and Agriculture” (funding agency: Bio-oriented Technology Research Advancement Institution, NARO). LG also acknowledges support from the European Centre of Excellence MaX (project ID 676598).

## Supplementary Materials

### Materials and Methods

#### Fluorescence-based assay of laccase-mediated detoxification of aflatoxin B_1_ and aflatoxin G2

Laccase from *Trametes versicolor* (Merck CAS80498) was dissolved in acetate buffer (pH 6.5) at a final concentration of 25 U/mL. Aflatoxin B_1_ and Aflatoxin G_2_ (Cayman Chemicals) were dissolved in LS-MS grade methanol (Merck) at 4 different concentrations: 3, 30, 50, and 100 μg/mL. Buffer solutions of laccase and aflatoxins were incubated at 28 °C over 96 hours under fast continuous shaking regime in a Synergy™ Mx Multi-Mode Microplate Reader (Biotek), each condition was performed in triplicate. Due to their natural fluorescence, aflatoxin concentration was fluorimetrically assayed (ex. 380 nm – em. 440 nm; gain 65 and 50 for AFB_1_ and AFG_2_, respectively); readouts were acquired every 10 minutes, totaling 577 data points by the end of the experiment. Controls were used assaying laccase fluorescence in the buffer in the absence of aflatoxins, and AFB_1_ and AFG_2_ fluorescence in the absence of laccase. To convert the fluorescence readout to the corresponding toxin concentration, a calibration curve was used based on measurements of a set of known toxin concentrations (Fig 1-FS1).

#### Mathematical modeling of laccase-aflatoxin interactions

The reaction kinetics were simulated using Matlab. The source codes are available on GitHub at: github.com/bmomeni/laccase-aflatoxins-reaction-kinetics.

#### Data analysis

The method of least squares (`lsqnonlin` in Matlab) was employed to fit the Michaelis-Menten kinetics to the experimental data. Detoxification rates were estimated by fitting a line to data from the first 100 minutes of the experiments for each initial toxin concentration.

#### Docking of aflatoxins with laccase

The 3D crystallographic structure of *Trametes versicolor* beta isoform laccase was retrieved from the Protein DataBank, accession code 1KYA [37]. The delta and gamma isoforms were derived instead via homology modeling using the beta isoform as a template as previously described [15]. The 3D structure of the aflatoxins were downloaded from PubChem (https://pubchem.ncbi.nlm.nih.gov/) [38], [39]. Before docking analysis, the consistency of atoms and bonds type for proteins and ligand were checked using the software Sybyl, version 8.1 (Certara USA, Princeton, NJ, USA), in agreement with a previous study [15]. Docking simulations were performed using the Protein Ligand Docking software GOLD (https://www.ccdc.cam.ac.uk/solutions/csd-discovery/components/gold/) and each docking pose was rescored using the HINT scoring function [40] for a better evaluation of the protein-ligand interaction, as previously reported [15].

##### Preparation of Protein and Ligand Structures for Docking

Protein homology models were cleaned up using the pdb4amber script, part of the Amber 2020 software suite [41]. Once structures passed inspection by Amber’s LEaP program, they were protonated using the H++ webserver version 3.2 [42]–[44] for a target pH of 6.5 to reflect real-use conditions. Because H++ does not account for metals, the resultant protonated structures were manually cleaned to flip histidine residues in order to maintain proper ligation of the embedded copper atoms. No explicit solvent molecules were included in the final structures. Ligand 3D structures were generated from ChemDraw 19.1 and optimized with Gaussian16 [45] in gas phase using HF/6-31G* basis set and functional. The resulting geometries were imported in Hermes, an application component of the CSD-Discovery Suite 2020 which interfaces with GOLD [46]. The bonds were repaired using Hermes’ structure clean up to ensure they were readable by GOLD.

##### Docking with GOLD

The protonated and adjusted protein structures were imported into the GOLD 2020.2.0 docking setup wizard. The pocket was defined using an atom from a residue lining the cavity, and GOLD’s pocket finding algorithm was used to then determine the pocket. Care was taken to ensure all residues and the copper atoms were recognized appropriately. All ligand flexibility options were enabled in addition to diverse solutions with a cluster size of 5 and RMSD of 2 Å. The genetic algorithm was set to the maximum search efficiency with automatic settings for the algorithm itself. Poses were scored and rescored with CHEMPLP and GoldScore respectively. The docking input files are available in the supplementary materials.

##### Rescoring with R-NiB

After docking was performed, the geometry was relaxed using the GFN-FF forcefield [47] as implemented in the XTB program with implicit solvation. The poses selected for further study were stripped of lone pairs using Hermes’ structure clean up. Atomic charges were calculated using Amber’s Antechamber program using the AM1-BCC charge model. PANTHER-0.18.21 [48] was used to generate negative images based on input files from Kurkinen et al, 2019 [31]. The resultant negative image Mol2 file was used as the query molecule for ShaEP [49] and the ligands were screened against it with the `- noOptimization` flag to use ShaEP as a rescoring function.

#### Quantum Mechanics-based modeling of a single electron oxidation effect on aflatoxin molecules

QM calculation were performed within the framework of Kohn-Sham Density Functional Theory (KS-DFT) [50], employing the Perdew–Burke-Ernzerhof (PBE) [51] exchange and correlation level of theory. The numerical results were extracted employing the BigDFT code [52], which uses Daubechies wavelets to express the KS orbitals. Hartwigsen-Goedecker-Hutter (HGH) pseudopotentials [53] were used to remove the core electrons orbitals. The use of wavelet basis sets enables one to control the precision of the results within a systematic approach, and at the same time to explicitly consider calculations of charged systems, as isolated boundary conditions are explicitly included in the calculations, without supercell aliasing effects, using the Poisson Solver of the code [54]. A wavelet grid spacing of 0.37 atomic units has been employed for the calculations presented in this work.

The code was used to calculate charged Delta-SCF, and the Fukui functions (FF) are defined as the difference between the neutral ground state electronic density and the corresponding quantity in the (vertical) cationic state. Such a definition of the FF is known to be more reliable—especially in the case of semilocal functionals—than the frontier orbital approach based on Koopmans theorem (see e.g. Ref. [42]).

To calculate the binding site, we performed KS-DFT calculations on the docked enzyme-toxin system using the linear scaling mode of the BigDFT code [27], [28] with the PBE approximation, HGH pseudopotentials, and a grid spacing of 0.4 atomic units. The charge of the enzyme was determined by minimizing the energy of the gas phase enzyme with respect to the number of electrons. Interaction strengths between system fragments were determined using the Fragment Bond Order tool as described in our previous study [23].

## Supplementary Data

### Calibration curves for fluorescence-based aflatoxins assay

Different starting concentrations of aflatoxin B_1_ and aflatoxin G_2_ (Cayman Chemicals) in acetate buffer (100 mM, pH 6.5) were employed to develop calibration curves to correlate measured fluorescence (ex. 380 nm; em. 440 nm) to toxin concentration. A Synergy™ Mx Multi-Mode Microplate Reader (Biotek) was used to perform the measurements.

**Fig 1-FS1.**
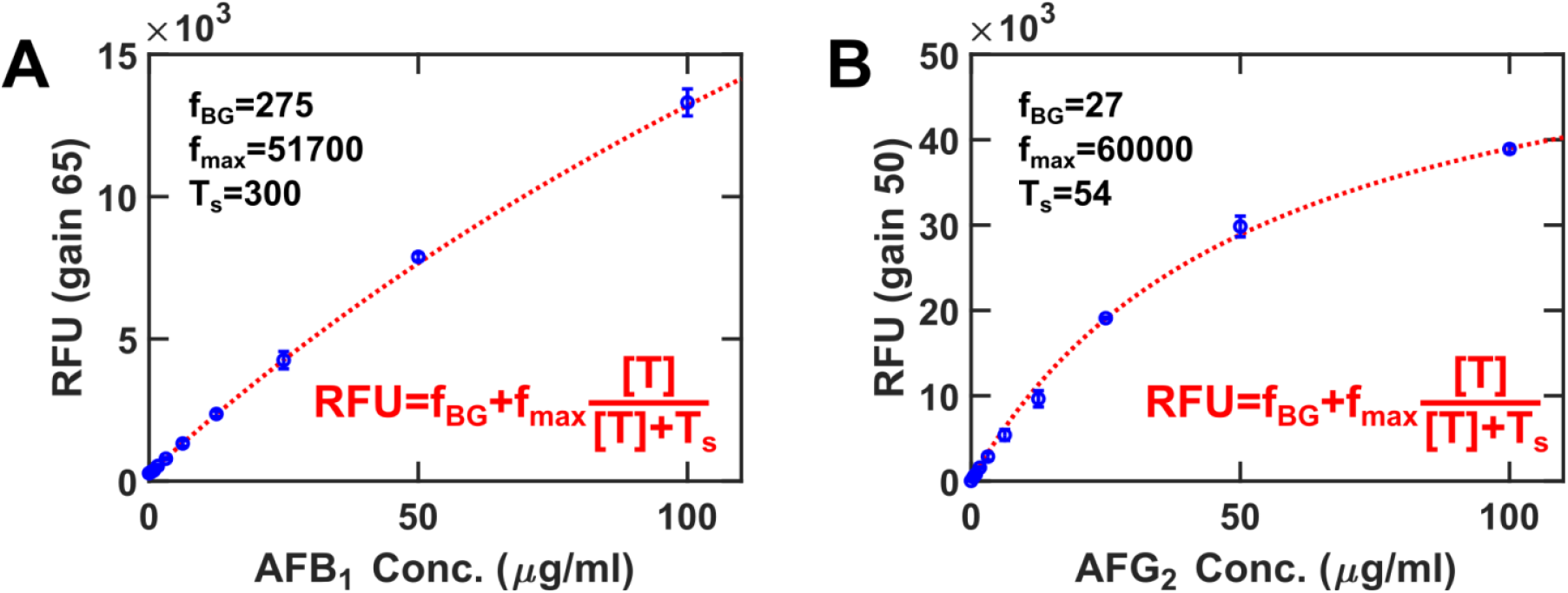
Calibration curves for fluorescence-assayed toxin concentration show reliable conversion between fluorescence and aflatoxin concentration. T represents toxin concentration; RFU represents Relative Fluorescence Units; T_s_, f_BG_ and f_max_ are fitting parameters; the fitting method is using the method of least squares as implemented in Matlab.

### Identification of degradation products of laccase activity on aflatoxins via LC/MS

50 U/mL laccase from *Trametes versicolor* (Sigma-Aldrich CAS80498) were added to 10 μg/mL toxin, Aflatoxin B_1_ and Aflatoxin G_2_ (Cayman Chemicals) separately, in acetate buffer (100 mM, pH 6.5) and left at 28 °C for 24 hrs. Degradation products were assayed under the following conditions:

#### LC/MS conditions

**Column:** Kinetex 2.6 μm EVO C18; 100 x 2.1 mm.
**Mobile phase A:** Water 5 mM Ammonium Acetate, 0.5% Acetic Acid
**Mobile phase B:** Methanol 5 mM Ammonium Acetate, 0.5% Acetic Acid
**Flow rate:** 350 μl/min
**UV Wavelength:** 354, 360 nm

| Time (min) | %A | %B |
| --- | --- | --- |
| Initial | 90 | 10 |
| 3 | 90 | 10 |
| 10 | 30 | 70 |
| 10.1 | 10 | 90 |
| 12 | 10 | 90 |
| 12.1 | 90 | 10 |
| 15 | 90 | 10 |

The eluent from the column was directed into the electrospray source of an Agilent 6220 TOF mass spectrometer operated in positive ionization mode. Data was converted into the mzML file format and analyzed using the MZMine software.

**Fig 2-FS1.**
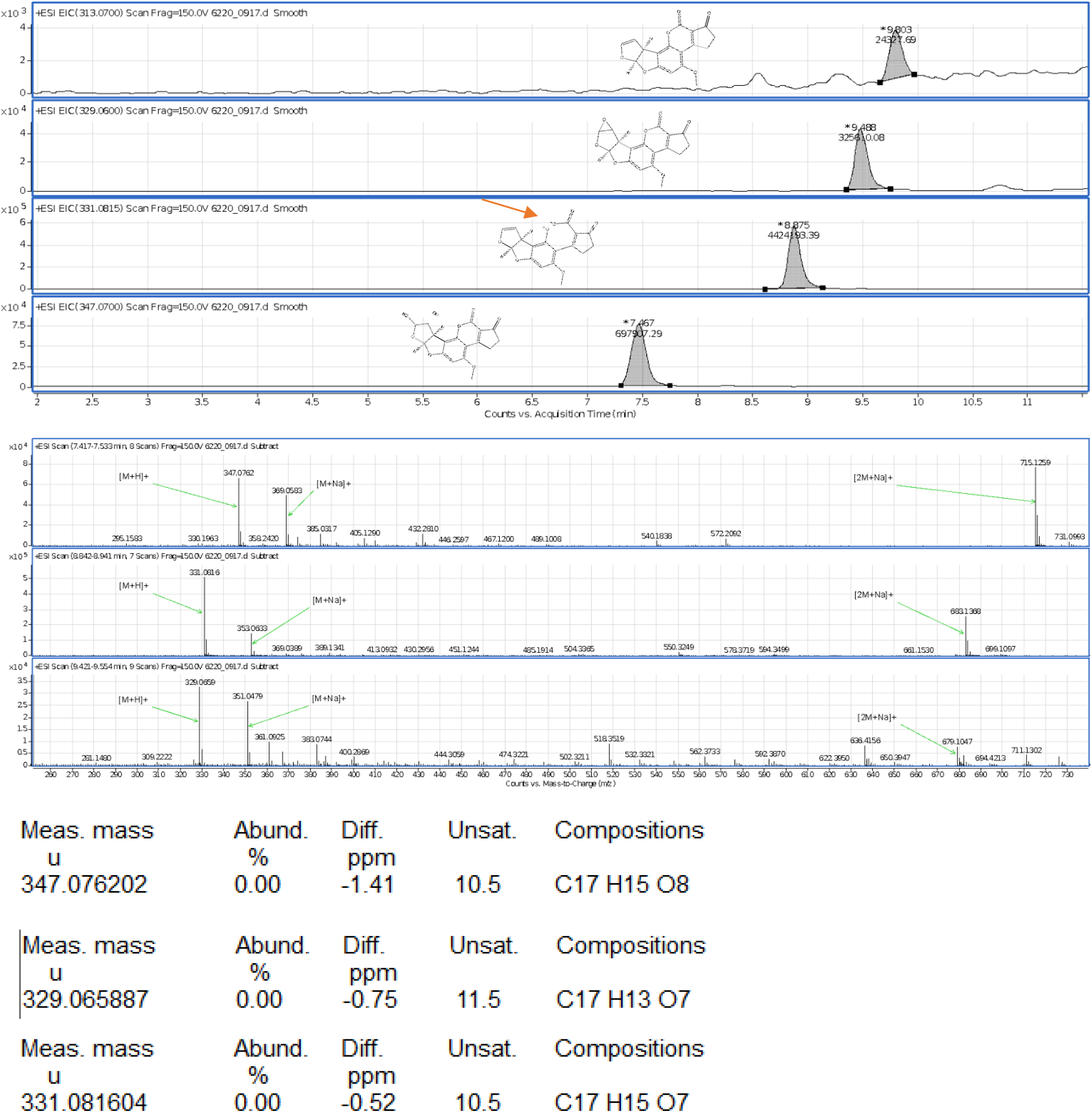
Mass spectroscopy reveals byproducts of AFB_1_ detoxification. The fully detoxified AFB_1_ molecule is the one with an open lactone ring, highlighted by the red arrow.

**Fig 2-FS2.**
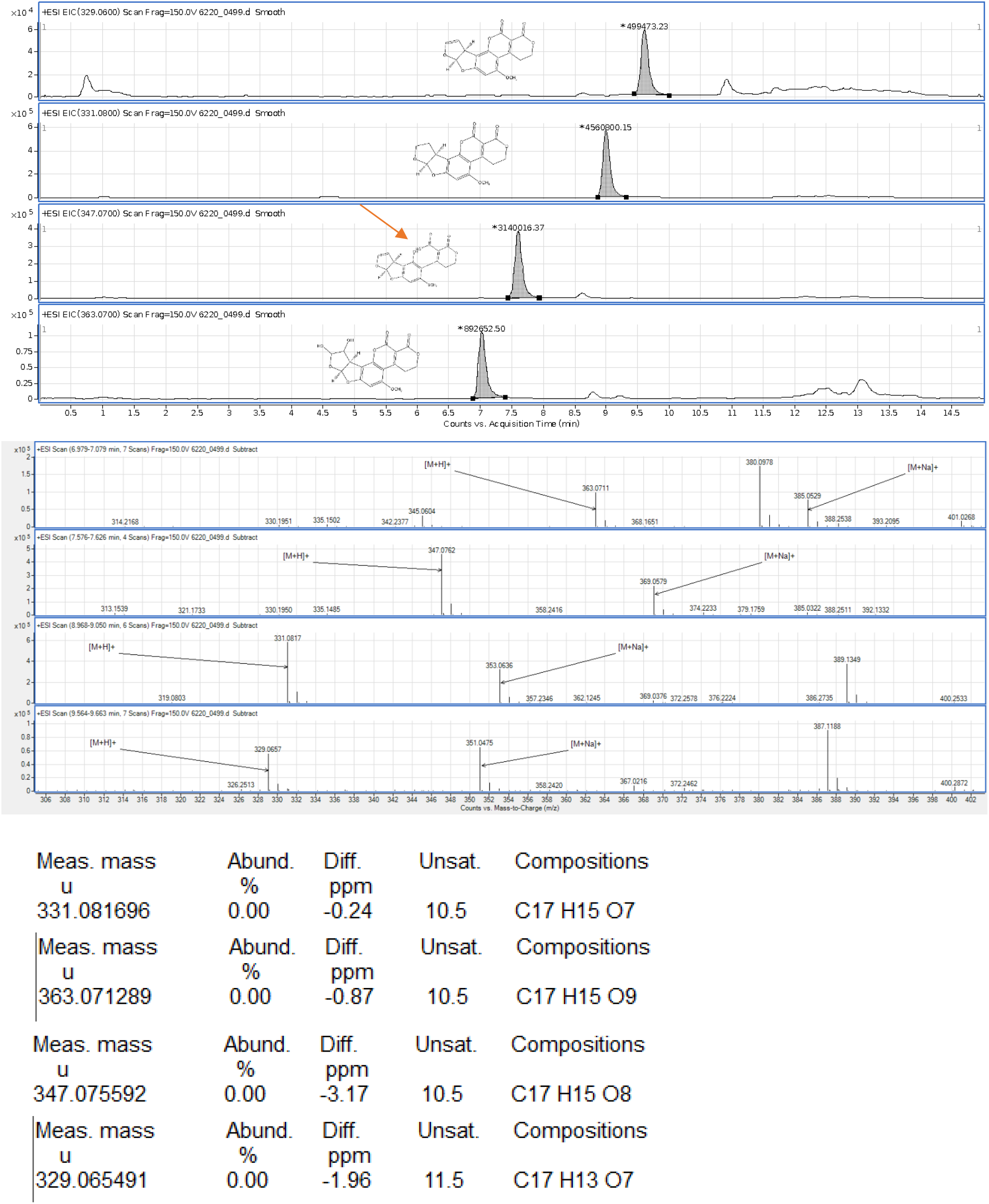
Mass spectroscopy reveals byproducts of AFG_2_ detoxification. The fully detoxified AFG_2_ molecule is the one with an open lactone ring, highlighted by the red arrow.

### R-NiB Rescoring of GOLD Docking Predictions

**Table S1.** R-NiB rescoring of poses presented in the paper. ShaEP considers both the geometric similarity of the molecule to the negative image and the similarity of the electronic field. The average of the two scores is the combined similarity.

| Molecule | Combined Similarity | Shape Similarity | Electronic Similarity |
| --- | --- | --- | --- |
| ABTS 15 | 0.0888454 | 0.0751505 | 0.10254 |
| ABTS 14 | 0.084475 | 0.0691252 | 0.0998248 |
| ABTS 9 | 0.0826305 | 0.0608398 | 0.104421 |
| ABTS 1 | 0.0800609 | 0.0583251 | 0.101797 |
| ABTS 6 | 0.0798537 | 0.0564544 | 0.103253 |
| G2 11 | 0.0752786 | 0.0448903 | 0.105667 |
| G2 13 | 0.075092 | 0.0475219 | 0.102662 |
| G2 1 | 0.0748488 | 0.0439506 | 0.105747 |
| B1 6 | 0.0743512 | 0.0469647 | 0.101738 |
| B1 2 | 0.0740255 | 0.0441423 | 0.103909 |
| G2 10 | 0.0738501 | 0.0435752 | 0.104125 |
| B1 15 | 0.0737891 | 0.0435919 | 0.103986 |
| G2 5 | 0.072414 | 0.0401788 | 0.104649 |
| B1 12 | 0.071184 | 0.0360877 | 0.10628 |

The results of R-NiB rescoring show major inversion in the rankings of the molecules where, for example, ABTS pose 15 had the lowest (worst) GOLDScore but is predicted the highest (best) by R-NiB.

**Fig 4-FS1.**
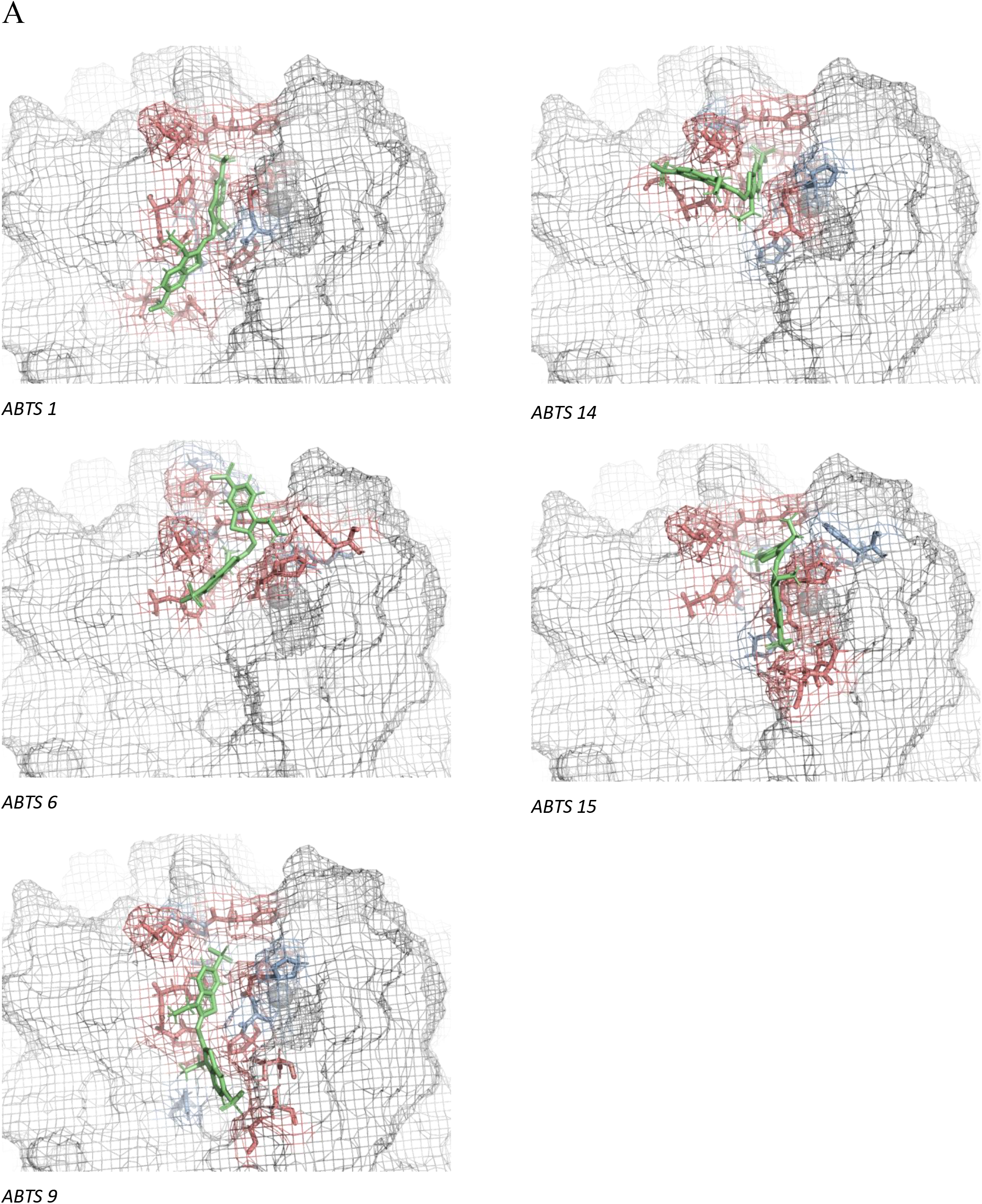
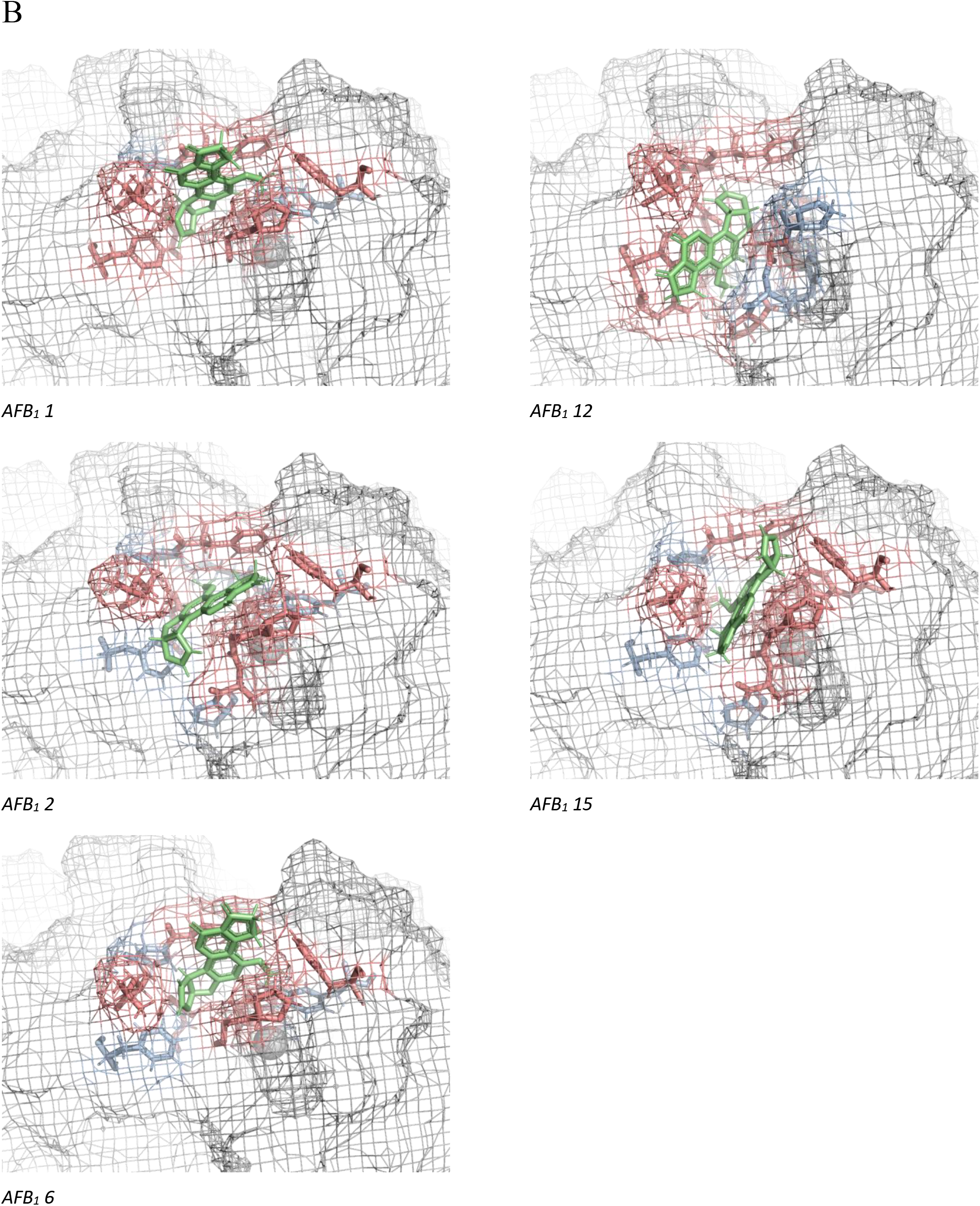
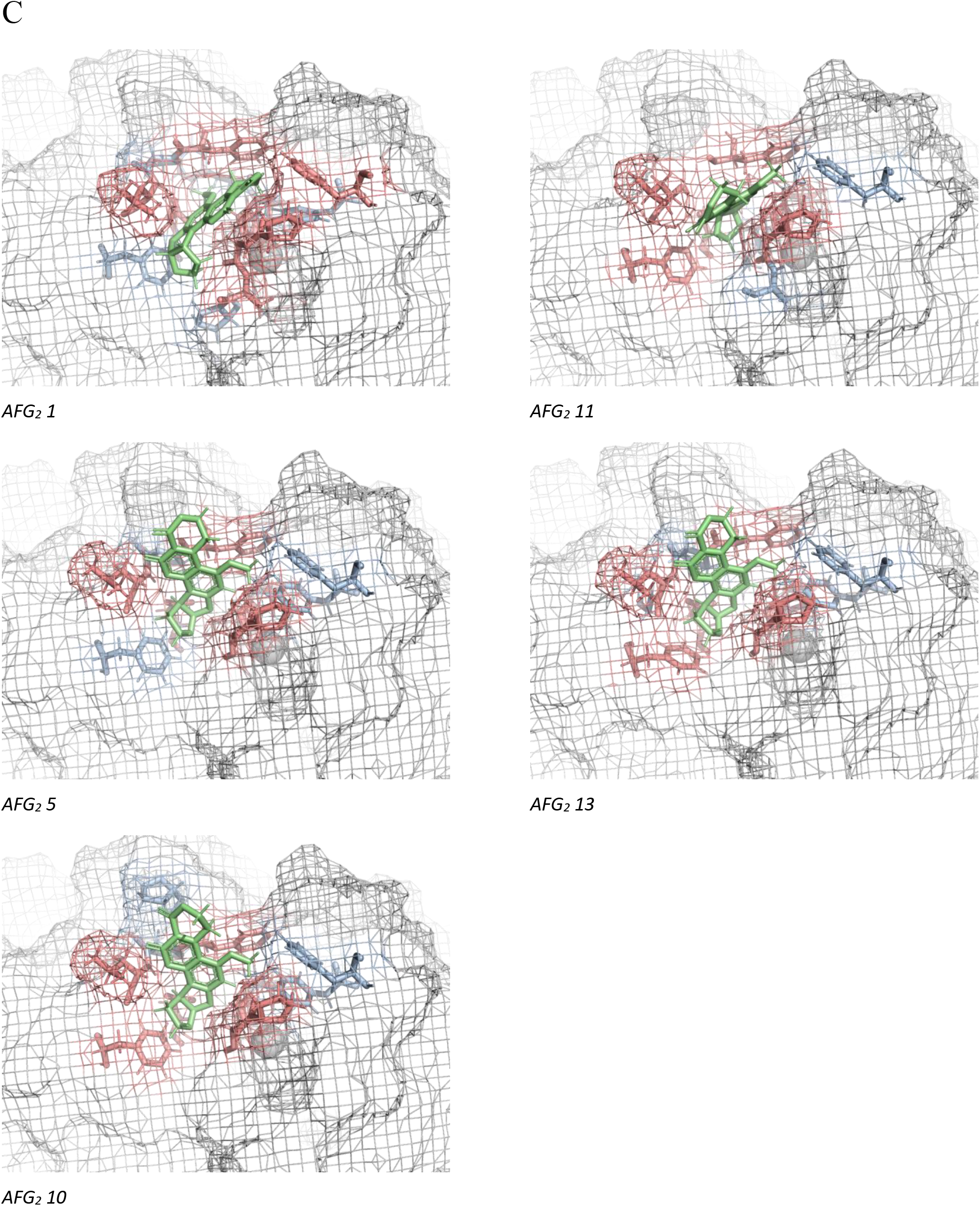
Representative images of the substrate specific active site of laccase are shown. Five poses are shown with (A) ABTS, (B) AFB_1_, and (C) AFG_2_ as the substrate.

## References

[1] H. J. Kulik, J. Zhang, J. P. Klinman, and T. J. Martínez, “How large should the QM region be in QM/MM calculations? the case of catechol O-methyltransferase,” J. Phys. Chem. B, vol. 120, no. 44, pp. 11381–11394, 2016, doi: 10.1021/acsjpcb.6b07814.

[2] L. E. Ratcliff et al., “Flexibilities of Wavelets as a Computational Basis Set for Large-Scale Electronic Structure Calculations,” pp. 1–32.

[3] R. Z. Liao and W. Thiel, “Convergence in the QM-only and QM/MM modeling of enzymatic reactions: A case study for acetylene hydratase,” J. Comput. Chem., vol. 34, no. 27, pp. 2389–2397, 2013, doi: 10.1002/jcc.23403.

[4] H. M. Senn and W. Thiel, “QM/MM methods for biomolecular systems,” Angew. Chemie - Int. Ed., vol. 48, no. 7, pp. 1198–1229, 2009, doi: 10.1002/anie.200802019.

[5] D. M. Mate and M. Alcalde, “Laccase: a multi-purpose biocatalyst at the forefront of biotechnology,” Microb. Biotechnol., vol. 10, no. 6, pp. 1457–1467, 2017, doi: 10.1111/1751-7915.12422.

[6] M. Zaccaria et al., “Designing a bioremediator: mechanistic models guide cellular and molecular specialization,” Curr. Opin. Biotechnol., vol. 62, pp. 98–105, 2020, doi: 10.1016/j.copbio.2019.09.006.

[7] E. I. Solomon, U. M. Sundaram, and T. E. Machonkin, “Multicopper oxidases and oxygenases,” Chem. Rev., vol. 96, no. 7, pp. 2563–2605, 1996, doi: 10.1021/cr950046o.

[8] M. Alcalde, “Laccases: biological functions, molecular structure and industrial applications.,” in Industrial Enzymes: Structure, Function and Applications, J. Polaina and A. P. MacCabe, Eds. Springer, 2007, pp. 461–478.

[9] J. W. Bennett and M. Klich, “Mycotoxins.,” Clin. Microbiol. Rev., vol. 16, no. 3, pp. 497–516, Jul. 2003, doi: 10.1128/cmr.16.3.497-516.2003.

[10] F. S. Chu, “Toxicology,” in Encyclopedia of Food Sciences and Nutrition, 2nd ed., no. April 2003, B. Caballero, L. C. Trugo, and P. M. Finglas, Eds. Elsevier Science, 2003, pp. 4096–4108.

[11] D. Klingelhöfer, Y. Zhu, M. Braun, M. H. K. Bendels, D. Brüggmann, and D. A. Groneberg, “Aflatoxin – Publication analysis of a global health threat,” Food Control, vol. 89, pp. 280–290, 2018, doi: 10.1016/j.foodcont.2018.02.017.

[12] I. Lyagin and E. Efremenko, “Enzymes for Detoxification of Various Mycotoxins: Origins and Mechanisms of Catalytic Action.,” Molecules, vol. 24, no. 13, p. 2362, Jun. 2019, doi: 10.3390/molecules24132362.

[13] Q. Wu, A. Jezkova, Z. Yuan, L. Pavlikova, V. Dohnal, and K. Kuca, “Biological degradation of aflatoxins,” Drug Metab. Rev., vol. 41, no. 1, pp. 1–7, Jan. 2009, doi: 10.1080/03602530802563850.

[14] J. F. Alberts, W. C. A. Gelderblom, A. Botha, and W. H. van Zyl, “Degradation of aflatoxin B1 by fungal laccase enzymes,” Int. J. Food Microbiol., vol. 135, no. 1, pp. 47–52, 2009, doi: 10.1016/j.ijfoodmicro.2009.07.022.

[15] L. Dellafiora, G. Galaverna, M. Reverberi, and C. Dall’Asta, “Degradation of aflatoxins by means of laccases from trametes versicolor: An in silico insight,” Toxins (Basel)., vol. 9, no. 1, 2017, doi: 10.3390/toxins9010017.

[16] M. Scarpari et al., “Aflatoxin control in maize by Trametes versicolor,” Toxins (Basel)., vol. 6, no. 12, pp. 3426–3437, 2014, doi: 10.3390/toxins6123426.

[17] R. Bourbonnais, M. G. Paice, I. D. Reid, P. Lanthier, and M. Yaguchi, “Lignin Oxidation by Laccase Isozymes from,” vol. 61, no. 5, pp. 1876–1880, 1995.

[18] N. Hakulinen and J. Rouvinen, “Three-dimensional structures of laccases,” Cell. Mol. Life Sci., vol. 72, no. 5, pp. 857–868, 2015, doi: 10.1007/s00018-014-1827-5.

[19] S. Lee, L. Dunn, A. DeLucca, and A. Ciegler, “Role of lactone ring of aflatoxin B1 in toxicity and mutagenicity.,” Experientia, vol. 37, no. 2, pp. 16–17, 1981.

[20] D. L. Nelson and M. M. Cox, Lehninger Principles of Biochemistry, 7th ed. Macmillan Learning, 2017.

[21] H. Zeinvand-Lorestani, O. Sabzevari, N. Setayesh, M. Amini, A. Nili-Ahmadabadi, and M. A. Faramarzi, “Comparative study of in vitro prooxidative properties and genotoxicity induced by aflatoxin B1 and its laccase-mediated detoxification products,” Chemosphere, vol. 135, pp. 1–6, 2015, doi: 10.1016/j.chemosphere.2015.03.036.

[22] C. Johannes and A. Majcherczyk, “Laccase activity tests and laccase inhibitors,” J. Biotechnol., vol. 78, no. 2, pp. 193–199, 2000, doi: 10.1016/S0168-1656(00)00208-X.

[23] W. Dawson, S. Mohr, L. E. Ratcliff, T. Nakajima, and L. Genovese, “Complexity Reduction in Density Functional Theory Calculations of Large Systems: System Partitioning and Fragment Embedding,” Journal of Chemical Theory and Computation 16, no. 5 (2020): 2952–2964.

[24] N. J. Christensen and K. P. Kepp, “Setting the stage for electron transfer: Molecular basis of ABTS-binding to four laccases from Trametes versicolor at variable pH and protein oxidation state,” J. Mol. Catal. B Enzym., vol. 100, pp. 68–77, 2014, doi: 10.1016/j.molcatb.2013.11.017.

[25] R. Mehra, J. Muschiol, A. S. Meyer, and K. P. Kepp, “A structural-chemical explanation of fungal laccase activity,” Sci. Rep., vol. 8, no. 1, pp. 1–16, 2018, doi: 10.1038/s41598-018-35633-8.

[26] L. E. Ratcliff, S. Mohr, G. Huhs, T. Deutsch, M. Masella, and L. Genovese, “Challenges in large scale quantum mechanical calculations,” Wiley Interdiscip. Rev. Comput. Mol. Sci., vol. 7, no. 1, pp. 1–18, 2017, doi: 10.1002/wcms.1290.

[27] “FDA Alert: Certain Lots of Pet Food from Multiple Brands Recalled for Aflatoxin | FDA.”.

[28] M. Awasthi, N. Jasiwal, S. Singh, V. P. Pandey, and U. N. Dwivedi, “Molecular docking and dynamics simulation analyses unraveling the differential enzymatic catalysis by plant and fungal laccases with respect to lignin biosynthesis and degradation.,” J. Biomol. Struct. Dyn., vol. 33, no. 9, pp. 1835–1849, 2014, doi: 10.1177/08959374970110010301.

[29] C. Martínez-Sotres, J. G. Rutiaga-Quiñones, R. Herrera-Bucio, M. Gallo, and P. López-Albarrán, “Molecular docking insights into the inhibition of laccase activity by medicarpin,” Wood Sci. Technol., vol. 49, no. 4, pp. 857–868, 2015, doi: 10.1007/s00226-015-0734-8.

[30] S. K. Kadam, A. S. Tamboli, S. B. Sambhare, B. H. Jeon, and S. P. Govindwar, “Enzymatic analysis, structural study and molecular docking of laccase and catalase from B. subtilis SK1 after textile dye exposure,” Ecol. Inform., vol. 48, no. October, pp. 269–280, 2018, doi: 10.1016/j.ecoinf.2018.10.003.

[31] S. T. Kurkinen, S. Lätti, O. T. Pentikäinen, and P. A. Postila, “Getting Docking into Shape Using Negative Image-Based Rescoring,” J. Chem. Inf. Model., vol. 59, no. 8, pp. 3584–3599, 2019, doi: 10.1021/acs.jcim.9b00383.

[32] G. Santiago et al., “Computer-Aided Laccase Engineering: Toward Biological Oxidation of Arylamines,” ACS Catal., vol. 6, no. 8, pp. 5415–5423, 2016, doi: 10.1021/acscatal.6b01460.

[33] E. Monza, M. F. Lucas, S. Camarero, L. C. Alejaldre, A. T. Martínez, and V. Guallar, “Insights into laccase engineering from molecular simulations: Toward a binding-focused strategy,” J. Phys. Chem. Lett., vol. 6, no. 8, pp. 1447–1453, 2015, doi: 10.1021/acs.jpclett.5b00225.

[34] S. Mohr et al., “Accurate and efficient linear scaling DFT calculations with universal applicability,” Phys. Chem. Chem. Phys., vol. 17, no. 47, pp. 31360–31370, 2015, doi: 10.1039/c5cp00437c.

[35] S. Mohr et al., “Daubechies wavelets for linear scaling density functional theory,” J. Chem. Phys., vol. 140, no. 20, 2014, doi: 10.1063/1.4871876.

[36] S. Mohr, M. Masella, L. E. Ratcliff, and L. Genovese, “Complexity Reduction in Large Quantum Systems: Fragment Identification and Population Analysis via a Local Optimized Minimal Basis,” J. Chem. Theory Comput., vol. 13, no. 9, pp. 4079–4088, 2017, doi: 10.1021/acs.jctc.7b00291.

[37] T. Bertrand et al., “Crystal structure of a four-copper laccase complexed with an arylamine: Insights into substrate recognition and correlation with kinetics,” Biochemistry, vol. 41, no. 23, pp. 7325–7333, 2002, doi: 10.1021/bi0201318.

[38] D. R. Lovley, “The microbe electric: conversion of organic matter to electricity,” Curr. Opin. Biotechnol., vol. 19, no. 6, pp. 564–571, 2008, doi: 10.1016/j.copbio.2008.10.005.

[39] S. Kim et al., “PubChem 2019 update: Improved access to chemical data,” Nucleic Acids Res., vol. 47, no. D1, pp. D1102–D1109, 2019, doi: 10.1093/nar/gky1033.

[40] G. Eugene Kellogg and D. J. Abraham, “Hydrophobicity: Is LogP(o/w) more than the sum of its parts?,” Eur. J. Med. Chem., vol. 35, no. 7–8, pp. 651–661, 2000, doi: 10.1016/S0223-5234(00)00167-7.

[41] D. M. Y. and P. A. K. D.A. Case, K. Belfon, I.Y. Ben-Shalom, S.R. Brozell, D.S. Cerutti, T.E. Cheatham, III, V.W.D. Cruzeiro, T.A. Darden, R.E. Duke, G. Giambasu, M.K. Gilson, H. Gohlke, A.W. Goetz, R. Harris, S. Izadi, S.A. Izmailov, K. Kasavajhala, A. Kovalenko, R. Krasny, T, “AMBER 2020.” 2020.

[42] R. Anandakrishnan, B. Aguilar, and A. V. Onufriev, “H++ 3.0: Automating pK prediction and the preparation of biomolecular structures for atomistic molecular modeling and simulations,” Nucleic Acids Res., vol. 40, no. W1, Jul. 2012, doi: 10.1093/nar/gks375.

[43] J. C. Gordon, J. B. Myers, T. Folta, V. Shoja, L. S. Heath, and A. Onufriev, “H++: A server for estimating pKas and adding missing hydrogens to macromolecules,” Nucleic Acids Res., vol. 33, no. SUPPL. 2, Jul. 2005, doi: 10.1093/nar/gki464.

[44] J. Myers, G. Grothaus, S. Narayanan, and A. Onufriev, “A simple clustering algorithm can be accurate enough for use in calculations of pKs in macromolecules,” Proteins Struct. Funct. Genet., vol. 63, no. 4, pp. 928–938, Jun. 2006, doi: 10.1002/prot.20922.

[45] M. J. Frisch et al., “Gaussian 16 Revision A.03.” 2016.

[46] G. Jones, P. Willett, R. C. Glen, A. R. Leach, and R. Taylor, “Development and validation of a genetic algorithm for flexible docking,” J. Mol. Biol., vol. 267, no. 3, pp. 727–748, Apr. 1997, doi: 10.1006/jmbi.1996.0897.

[47] S. Spicher and S. Grimme, “Robust Atomistic Modeling of Materials, Organometallic, and Biochemical Systems,” Angew. Chemie Int. Ed., vol. 59, no. 36, pp. 15665–15673, Sep. 2020, doi: 10.1002/anie.202004239.

[48] S. P. Niinivehmas, K. Salokas, S. Lätti, H. Raunio, and O. T. Pentikäinen, “Ultrafast protein structure-based virtual screening with Panther,” J. Comput. Aided. Mol. Des., vol. 29, no. 10, pp. 989–1006, Oct. 2015, doi: 10.1007/s10822-015-9870-3.

[49] M. J. Vainio, J. S. Puranen, and M. S. Johnson, “ShaEP: Molecular overlay based on shape and electrostatic potential,” J. Chem. Inf. Model., vol. 49, no. 2, pp. 492–502, Feb. 2009, doi: 10.1021/ci800315d.

[50] W. Kohn and L. J. Sham, “Self-Consistent Equations Including Exchange and Correlation Effects*,” Phys. Rev., vol. 385, no. 1951, 1965, doi: 10.1103/PhysRev.140.A1133.

[51] J. P. Perdew, K. Burke, and M. Ernzerhof, “Generalized Gradient Approximation Made Simple,” Phys. Rev. Lett., vol. 77, no. 18, pp. 3865–3868, 1996, doi: 10.1103/PhysRevLett.77.3865.

[52] L. Genovese et al., “Daubechies wavelets as a basis set for density functional pseudopotential calculations,” J. Chem. Phys., vol. 129, no. 1, 2008, doi: 10.1063/1.2949547.

[53] A. Willand et al., “Norm-conserving pseudopotentials with chemical accuracy compared to all-electron calculations,” J. Chem. Phys., vol. 138, no. 10, 2013, doi: 10.1063/1.4793260.

[54] A. Cerioni, L. Genovese, A. Mirone, and V. A. Sole, “Efficient and accurate solver of the three-dimensional screened and unscreened Poissons equation with generic boundary conditions,” J. Chem. Phys., vol. 137, no. 13, 2012, doi: 10.1063/1.4755349.

